# Trans-synaptic Fish-lips Signaling Prevents Misconnections between Non-synaptic Partner Olfactory Neurons

**DOI:** 10.1101/600064

**Authors:** Qijing Xie, Bing Wu, Jiefu Li, Hongjie Li, David J Luginbuhl, Chuanyun Xu, Xin Wang, Liqun Luo

**Author notes:** Q.X. and B.W. contributed equally to the work.

## Abstract

Our understanding of the mechanisms of neural circuit assembly is far from complete. Identification of new wiring molecules with novel mechanisms of action will provide new insights into how complex and heterogeneous neural circuits assemble during development. Here, we performed an RNAi screen for cell-surface molecules and identified the leucine-rich-repeat containing transmembrane protein, Fish-lips (Fili), as a novel wiring molecule in the assembly of the *Drosophila* olfactory circuit. Fili contributes to the precise targeting of both olfactory receptor neuron (ORN) axons as well as projection neuron (PN) dendrites. Cell-type-specific expression and genetic analyses suggest that Fili sends a trans-synaptic repulsive signal to neurites of non-partner classes that prevent their targeting to inappropriate glomeruli in the antennal lobe.

**Significance Statement:** In the fruit fly olfactory system, 50 classes of olfactory receptor neurons (ORNs) make precise synaptic connections with 50 classes of corresponding projection neurons (PNs). Identification of wiring molecules in this circuit can provide insight into understanding neural circuit assembly. This paper reports the role of a transmembrane protein, Fish-lips (Fili), in forming specific connections in this circuit. We found that some ORN axons are repelled by Fili, which is present on dendrites of non-matching PN class, preventing them from targeting inappropriate glomeruli. Similarly, some PN dendrites are repelled by Fili expressed by non-matching ORN class for their correct targeting. Together, these results suggest that Fili mediates repulsion between axons and dendrites of non-synaptic partners to ensure precise wiring patterns.

## Introduction

The brain comprises of extremely complex yet precisely-wired circuits of neurons which allow animals to faithfully relay and process information. In order to establish these specific neural connections, a coordinated sequence of developmental steps is taken: axons and dendrites first navigate to their target zones, they then identify the appropriate partners, and finally form functional synapses. Secreted and membrane-bound factors are required for each of these steps (1–5). These factors can act as either ligands or receptors to allow a developing neurite to sample its environment and select the correct target. Although much progress has been made in both identifying wiring-related molecules and investigating cellular mechanisms underlying each of the steps described above, our understanding of neural circuit assembly is far from complete. Therefore, it is important to identify novel molecules and mechanisms that contribute the precise assembly of neural circuits.

To gain more insight into this developmental process, we studied the *Drosophila* olfactory system. Here, each of the 50 classes of olfactory receptor neurons (ORNs) extend their axons that make synaptic connections with dendrites of their corresponding projection neuron (PN) class in 50 anatomically discrete glomeruli in the antennal lobe (8–10). Because of the stereotyped organization of different synaptic partner pairs, the richness of connections, and the availability of genetic tools, many wiring molecules have been identified and studied in this circuit (11, 12). Furthermore, novel wiring molecules discovered in this system have guided our understanding of the wiring logic of other neural circuits across different species (13–16).

Here, we designed and carried out an RNAi screen to identify novel wiring molecules in a region of the antennal lobe that has not been extensively studied. From this screen, we identified a novel wiring molecule, Fish-lips (Fili), a leucine-rich repeat (LRR)-containing transmembrane protein previously known for its function in regulating dioxin receptor homolog Spineless-mediated apoptosis (17). We investigated the function of Fili in both ORN axon and PN dendrite targeting during olfactory circuit development. We found that Fili signals reciprocally between ORNs and PNs to instruct precise target selection.

## Results

### A novel leucine-rich repeat protein, Fish-lips, is required for the wiring of olfactory neurons

To identify new molecules underlying the *Drosophila* olfactory circuit wiring specificity, we have previously carried out genetic screens for ORN axon and PN dendrite targeting to the anterior-lateral side of the antennal lobe (18, 19). While certain molecules broadly contribute to the precise wiring of many types of olfactory neurons (20, 21), others have been found to be region- and even glomerulus-specific (6, 7, 11, 13, 18, 22). We reasoned that neurons in different locations in the antennal lobe might use a different set of wiring molecules for their circuit assembly. To identify new specific PN drivers with reliable expression in other regions of the antennal lobe, we screened through a collection of enhancer-GAL4 lines from the FlyLight project (23, 24). We found five lines that show robust, specific, and consistent labeling patterns of various PN subtypes (Fig. S1). The labeled glomeruli covered distinct regions including the dorsolateral, ventromedial, and middle regions of the antennal lobe. We also converted these enhancer-GAL4 lines into another binary system, LexA/LexAop, by fusing each of the specific enhancer fragments to the *LexA* sequence (25). Using one of the lines, *GMR86C10-LexA*, we designed a new screening scheme to identify novel wiring molecules that function in the ventromedial antennal lobe.

Specifically, we used *GMR86C10-LexA>LexAop-mtdT* to label dendrites of VM5v and VM5d PNs, and used *Or98a-mCD8-GFP* and *Or92a-rCD2* to label axons of VM5v and VA2 ORNs, respectively (Fig. 1 *A* and *B*). In addition to these neuronal-class specific markers, we visualized the neuropil using an antibody against N-cadherin. Using the pan-neuronal *C155-GAL4* driver line, we expressed RNAi against predicted transmembrane and secreted molecules. We tested around 700 RNAi lines covering over 200 genes whose protein products contain domain types of LRR, immunoglobulin, cadherin, epidermal growth factor repeat, and fibronectin for their necessity in olfactory neuron wiring accuracy.

**Figure 1.**
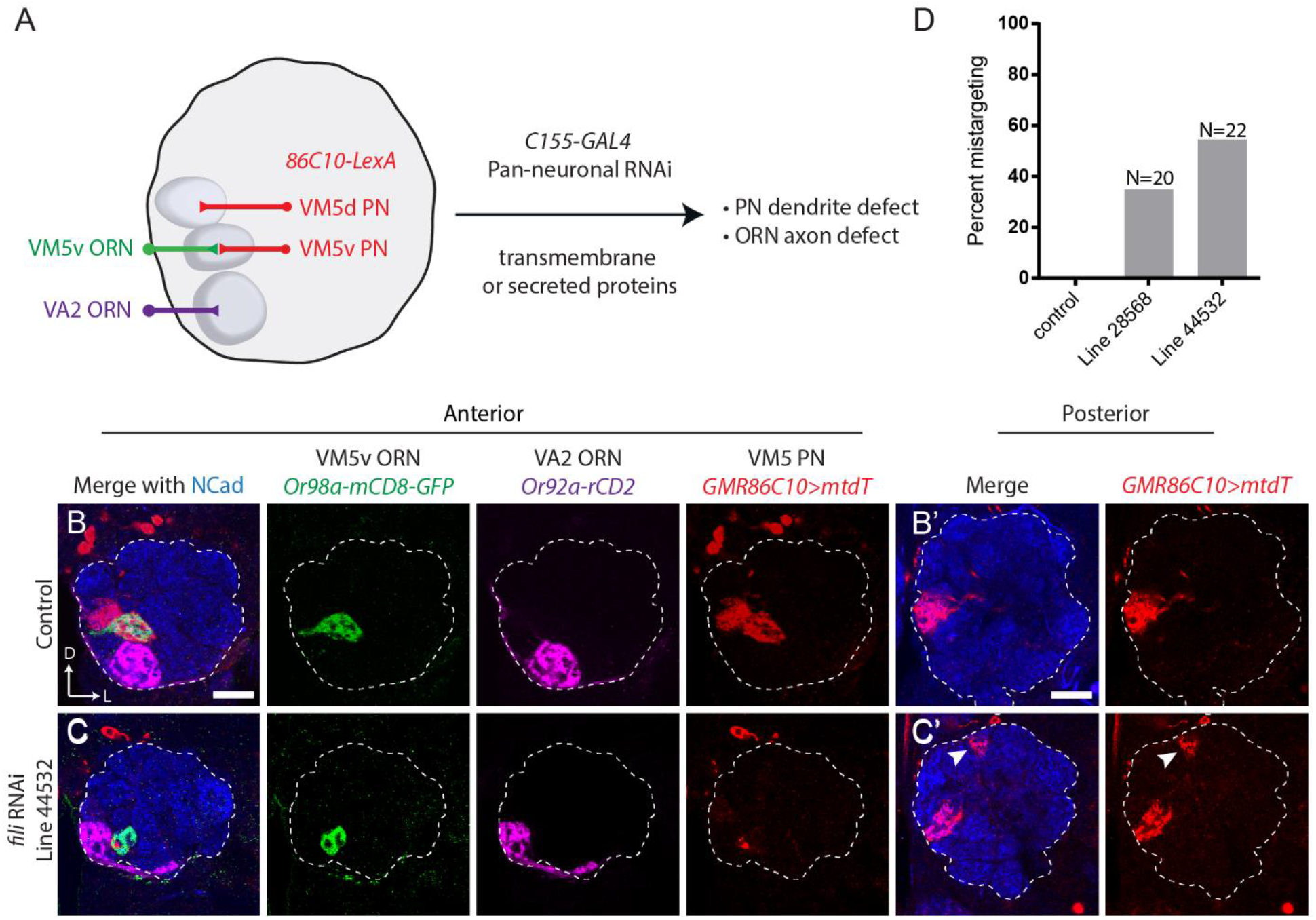
Identification of Fish-lips as a wiring specificity molecule through an RNAi screen. (*A*) Schematic of RNAi screen. Pan-neuronal *C155-GAL4* was used to drive *UAS-RNAi* against predicted transmembrane and secreted molecules. Dendrites of two PN classes, VM5v and VM5d (hereafter VM5 PNs), are labeled by *GMR86C10-LexA>LexAop-mtdT*. Axons of VM5v and VA2 ORNs are labeled by two different markers, *Or98a-mCD8GFP* and *Or92a-rCD2* respectively. (*B and C*) Targeting of dendrites of VM5 PNs (red) and axons of VM5v (green) and VA2 ORNs (magenta) in the antennal lobe on the anterior side. Neuropil staining by the Ncad antibody are shown in blue. Dashed line outlines the antennal lobe neuropil. (*B*’ and *C*’) Posterior sections of the same antennal lobes as in *B* and *C*. Dendrites of VM5 PNs and VM5v/VA2 ORN axons target to their corresponding glomeruli in control (*B*). *C155-GAL4* driven *UAS-fili-RNAi* (VDRC 44532) shows PN dendrite targeting defect (*C*). Ectopic PN targets are indicated by arrowheads on the posterior section of (*C*’). (*D*) Quantification of VM5 PNs mistargeting of two different RNAi lines against *fili* (Bloomington 28568 and VDRC 44532). The phenotype penetrance of two independent RNAi lines are 7/20 and 12/22, respectively. Scale bars, 20 μm. D, dorsal; L, lateral.

From the screen, we identified a candidate wiring specificity protein, Fili-lips (Fili). In wild-type animals, *GMR86C10-LexA>LexAop-mtdT* PNs extend their dendrites to only two glomeruli—VM5v and VM5d (hereafter VM5 PNs). When *fili* was knocked-down by either one of the two RNAi lines against different regions of *fili*, VM5 PN dendrites ectopically invaded a glomerulus more dorsal and posterior to the correct target (arrowhead in Fig. 1*C*’ and Fig. S2*A*), with penetrance correlated with *fili* knockdown efficiency (Fig. 1*D* and S2*B*).

*fili* encodes a transmembrane protein that contains 14 LRR domains (Fig. S3*A*). Fili shares highest amino acid sequence similarities with two other LRR proteins, Capricious (Caps) and Tartan (Trn) (17), both of which have been demonstrated to regulate PN dendrite targeting (22). However, the function of Fili in the nervous system has not been investigated previously.

To investigate the function of Fili, we generated a null allele using CRISPR-mediated gene editing, which removed the first exon and part of the second exon of *fili* (Fig. 2*A*). According to SMART protein prediction (26), this excision should eliminate the DNA sequence that encodes the start codon, the signal peptide, and more than half of the LRR domains. We tested this allele by staining brains of *fili*^−/−^ flies with a Fili antibody we generated against an epitope on the intracellular domain of Fili (Fig. S3*A*), and we could not detect any signal (Fig. 2*C*). Because the targeting epitope of Fili antibody is not contained in the deleted region, our data suggested that this deletion fully disrupted production of Fili. Similar to the phenotype we observed by pan-neuronal RNAi knockdown of *fili, fili*^−/−^ animals show dorsoposterior ectopic targeting of VM5 PN dendrites (Fig. 2*E*’ arrowhead).

**Figure 2.**
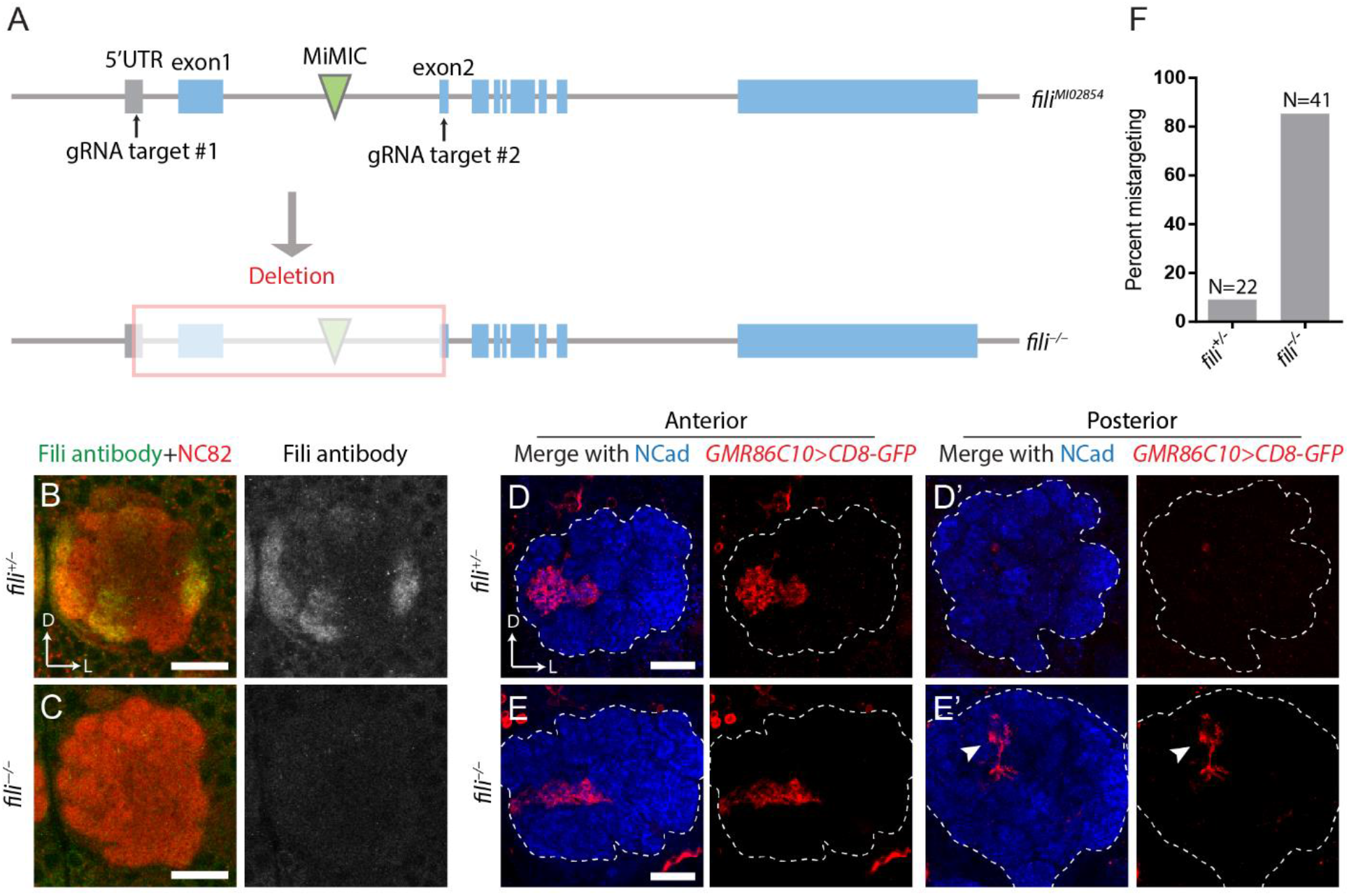
*fili* mutant recapitulates RNAi phenotype. (*A*) Schematic for the generation of *fili* null mutant by CRISPR-mediated excision. *vas-Cas9; fili^MI02854^* eggs were co-injected with two gRNAs targeting the 5’ UTR (untranslated region) and the second exon. The first coding exon and part of the second coding exon are deleted in the mutant. Blue bars: coding exons of *fili*. (*B and C*) Maximum projection of anti-Fili serum staining (green and grey) of heterozygous control (*B*) and *fili*^−/−^ fly (*C*). (*D and E*) Targeting of VM5 PNs labeled by *GMR86C10-GAL4>UAS-mCD8GFP* (shown in red) in the antennal lobe of heterozygous *fili*^+/−^ (*D*; mistargeting ratio: 2/22) or Homozygous *fili*^−/−^ animals (*E*; mistargeting ratio: 35/41). (*F*) Quantification of mistargeting ratio from panels (*D*) and (*E*). Scale bars, 20 μm.

### Fili is expressed in a sparse set of ORNs and PNs

The expression pattern of a wiring molecule can be informative for understanding its mechanism of action. While guidance molecules with a graded expression pattern are often important for the initial coarse targeting of developing neurites (27, 28), wiring molecules with discrete patterning are usually used to refine the final targeting to a specific region (22). In *fili*^−/−^ animals, VM5 PN dendrites consistently mistargeted to the same glomerulus. This observation suggests that Fili could serve as a discrete signal for VM5 PN dendrites during their final target selection and refinement.

To assay the expression of Fili, we stained brains at 48 hours after puparium formation (48h APF). At this time, matching between ORN axons and PN dendrites has just been established, and the expression pattern likely reflects the molecules used for the final targeting. Additionally, because discrete glomeruli have just formed at this stage, discerning the identity of neurons based on their projection pattern is possible. Immunostaining with anti-Fili antibodies revealed the presence of Fili protein in a sparse set of glomeruli in the developing antennal lobe (Fig. 3*A*).

**Figure 3.**
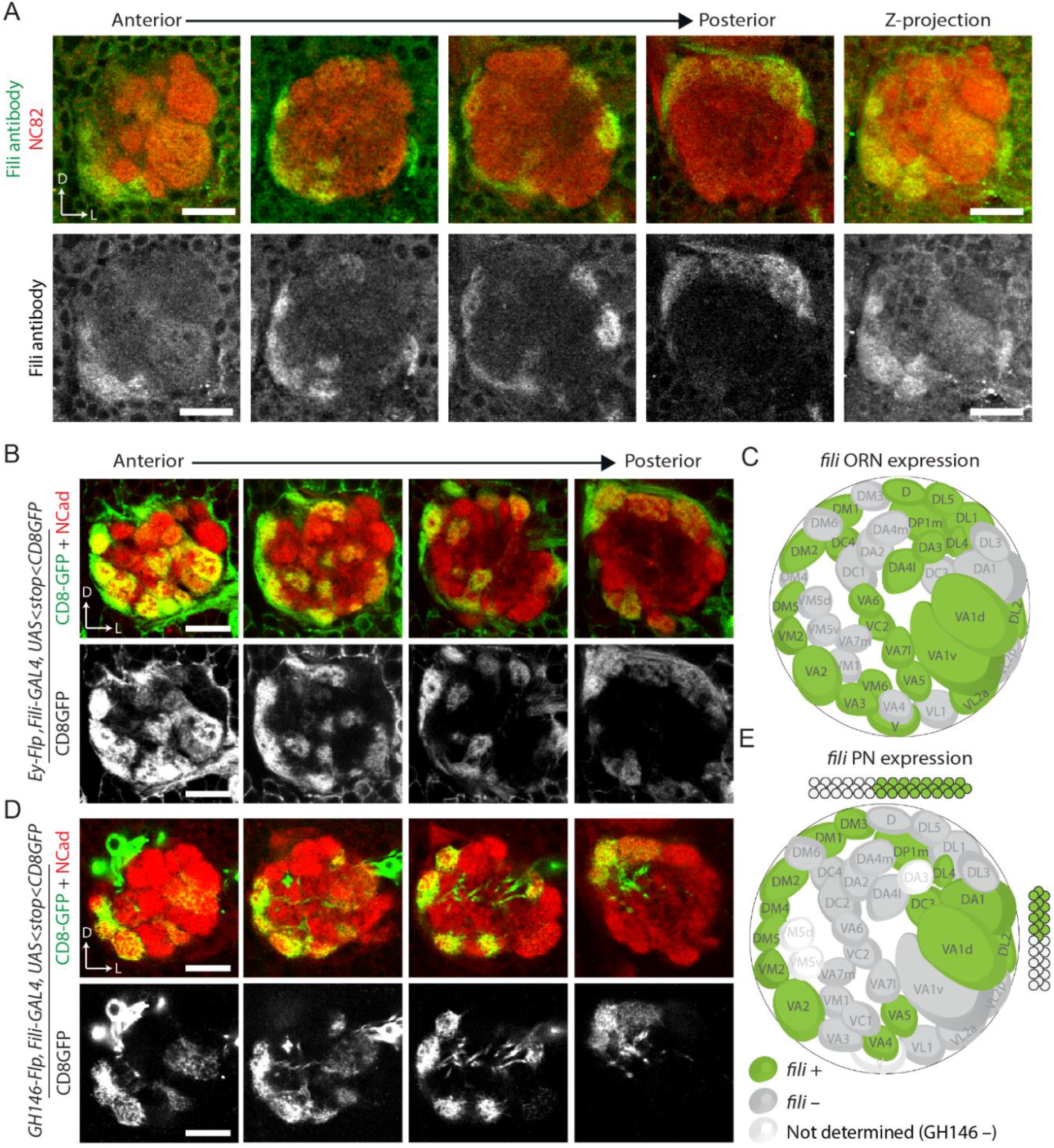
Fili is differentially expressed in a subset of ORNs and PNs during development. (*A*) anti-Fili serum staining of antennal lobe in wild-type animal (*w^1118^*) at 48 hours APF (green and grey). Neuropil is stained by the NC82 antibody (red). (*B and D*) *ey-FLP* or *GH146-FLP* intersecting with *fili-GAL4* using *UAS-FRT-stop-FRT-mCD8GFP* as a reporter shows *fili* expression pattern in ORNs (B) or PNs (D) at 48 hours APF. (*C and E*) Schematic 2-dimensional representation of the glomerular innervation pattern of *fili-GAL4-expressing* ORNs (*C*) or pNs (*E*). Scale bars, 20 μm.

Glomerular-specific Fili patterns could be due to its expression in ORNs, in PNs, or both. To distinguish these possibilities, we generated a transcriptional reporter *fili-GAL4* (29) so as to study its cell-type specific expression using cell type-specific Flp drivers and an intersectional reporter. An artificial exon containing a splicing acceptor, an in-frame *T2A-GAL4*, and a transcription terminator was inserted into a MiMIC locus (*MI02854*) residing in a coding intron of *fili* (Fig. S3*B*). To validate that this line faithfully represents Fili expression, we crossed it to a membrane fluorescent reporter (*UAS-mCD8-GFP*) and observed similar GFP pattern compared to Fili protein pattern in the antennal lobe (Fig. S3*C*). The mCD8-GFP reporter for *fili-GAL4* showed bright cell body labeling around the antennal lobe, but such pattern was not present in the staining of Fili antibody. This is likely because the mCD8-GFP reporter does not reflect sub-cellular localization of the Fili protein.

To visualize the contribution of Fili by ORNs and PNs, respectively, we intersected *fili-GAL4* with FLP recombinase lines expressed in either ORNs (*ey-FLP*) or PNs (*GH146-FLP*). When combined with *UAS-FRT-STOP-FRT-mCD8GFP*, mCD8-GFP would be solely produced in *fili* positive ORNs or PNs. Using this strategy, we observed that Fili was expressed sparsely in ORNs and PNs in a “salt and pepper”-like pattern (Fig. 3 *B* and *D*), reminiscent of the expression of Caps and Trn (22). This expression pattern suggests that Fili likely serves as a discrete determinant that constrains glomerular targeting of neurites, instead of setting a gradient for trajectory selection or coarse targeting. Further analysis revealed no statistically significant correlation of *fili* expression between ORNs and PNs (Fisher’s exact test, p-value = 0.4704). This suggest that Fili unlikely mediates homophilic attraction or repulsion between ORNs and PNs.

### Fili in ORNs signals to VM5 PNs for proper dendrite targeting

To determine which neurons require Fili functions for the proper targeting of VM5 PN dendrites, we carried out mosaic analyses with a repressible cell marker (MARCM) (30) to test a possible cell-autonomous function. Using hsFLP-based MARCM, we generated *fili*^−/−^ neuroblast clones and observed no mistargeting of VM5 PN dendrites as visualized by *GMR86C10-GAL4* (Fig. 4 *A* and *C*). This indicates that Fili is not required in VM5 PNs for their correct dendrite targeting. In contrast, deleting *fili* in most ORNs using *ey-FLP-* based MARCM combined with a cell-lethal strategy (31) recapitulated whole-animal *fili*^−/−^ phenotype (Fig. 4 *B* and *C*).

**Figure 4.**
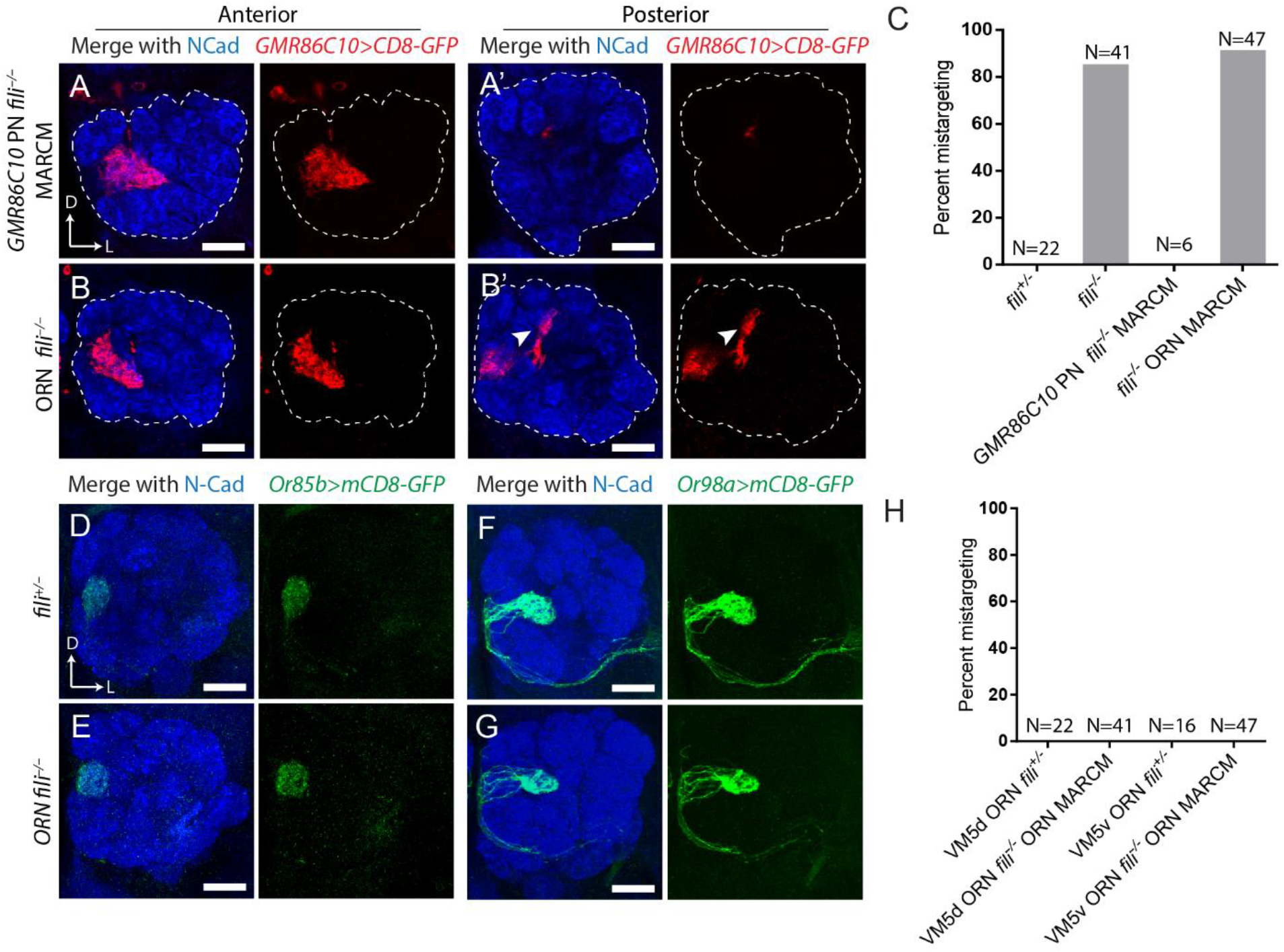
Fili in ORNs signals to VM5 PNs for the correct targeting of their dendrites. (*A*) Dendrite targeting of *fili*^−/−^ VM5 neuroblast clone produced by *hsFLP* MARCM in a *fili*^+/−^ background (mistargeting in 0/6 antennal lobes). (*B*) Dendrite targeting of *fili*^+/−^ VM5 PNs in animals with the majority of ORNs being *fili*^−/−^ generated by *ey-FLP* MARCM combined with cell-lethal strategy (mistargeting in 43/47 antennal lobes). (*A*’ and *B*’) Posterior sections of the same antennal lobe with arrowhead highlighting ectopic targeting of VM5 PN dendrites. (*C*) Quantification of *GMR86C10+* PN mistargeting from panel (*A*) and (*B*). (*D and E*) Targeting of VM5d ORN axons labeled by *Or85b-GAL4>UAS-mCD8GFP* in *fili*^+/−^ animals (mistargeting in 0/22 antennal lobes) (*D*) and in ORNs *fili*^−/−^ (mistargeting in 0/41 antennal lobes) (*E*). (*F* and *G*) Targeting of VM5v ORN axons labeled by *Or98a-GAL4>UAS-mCD8GFP* in the antennal lobe of *fili*^+/−^ animals (mistargeting in 0/16 antennal lobes) (*F*) and in ORNs *fili*^−/−^ animals (mistargeting in 0/47 antennal lobes) (*G*). (*H*) Quantification of mistargeting of VM5d and VM5v ORN axons from panel (*D-G*). Scale bars, 20 μm.

Two mechanisms might account for the above results. First, Fili is required in VM5 ORNs for their correct axon targeting. When those ORN axons mistarget, dendrites of their partner PN classes mistarget with them (18). Second, Fili in ORNs signal to dendrites of VM5 PNs to direct their correct targeting. To test the first possibility, we removed Fili in ORNs and visualized axon targeting of VM5v or VM5d ORNs. However, no obvious axon mistargeting was observed (Fig. 4*D-H*). This ruled out the first possibility and suggested that Fili expressed by ORNs to signal to VM5 PNs for their dendrite target selection. To understand where the Fili signal originates, we analyzed the expression pattern of Fili in ORNs. We observed no Fili expression in VM5v and VM5d ORNs (Fig. S4*A*) at 48h APF. By contrast, the mistargeted site had high Fili signal in ORNs (Fig. S4*B*). This observation suggests that Fili expressed in the ORN class occupying the ectopic target site repels VM5 PNs to prevent their dendrites from targeting to inappropriate glomeruli.

### Fili is required for the correct targeting of a small subset of PNs and ORNs

Because Fili is expressed in many ORN and PN classes, we examined its involvement in the wiring process of other neuronal classes. Identifying more classes of neurons that require Fili for their targeting would allow us to test whether Fili repels other neurites as a general mechanism of action.

We labeled different ORN and PN classes in *fili*^−/−^ background to investigate their axon or dendrite targeting. We examined dendritic targeting of 9 PN classes using 6 drivers, and did not observe any obvious targeting defects (Fig. S5). We also labeled 13 different ORN classes in *fili*^−/−^ background, and observed 3 classes with abnormal axon targeting patterns (Fig. S6). Among the three ORN classes that showed axon targeting abnormalities, VA7l ORNs displayed a misshaped glomerulus. However, we did not observe ectopic targeting of VA7l ORN axons, unlike the other two classes (Fig. S6*C*). This phenotype suggests that the VA7l ORN abnormality could be a secondary effect caused by PN mistargeting or its own mistargeting to neighboring glomeruli. For both DC1 ORNs and VA1v ORNs, we observed clear ectopic targeting of their axons (Fig. S6 *B* and *D*), indicating that Fili is likely required for their correct axon target specificity. We chose to focus on VA1v ORNs because there is a wealth of tools available that allows us to manipulate gene expression in regions adjacent to VA1v to investigate how Fili regulates wiring.

### Fili is required in VA1d/DC3 PNs to prevent ectopic targeting of VA1v ORN axons

In wild-type flies, ORNs of different classes target their axons to distinct glomeruli and never intermingle (Fig. 5*A*). However, in *fili*^−/−^ flies, VA1v ORN axons invaded the VA1d glomerulus along with VA1d axons (Fig. 5 *B* and *H*). When we removed *fili* in most ORNs by ey-FLP-based MARCM combined with cell lethal strategy, we did not observe any targeting defect of VA1v ORN axons (Fig. 5 *C* and *H*). Thus, Fili expressed in VA1v ORNs or other ORNs is not required for their axon targeting.

**Figure 5.**
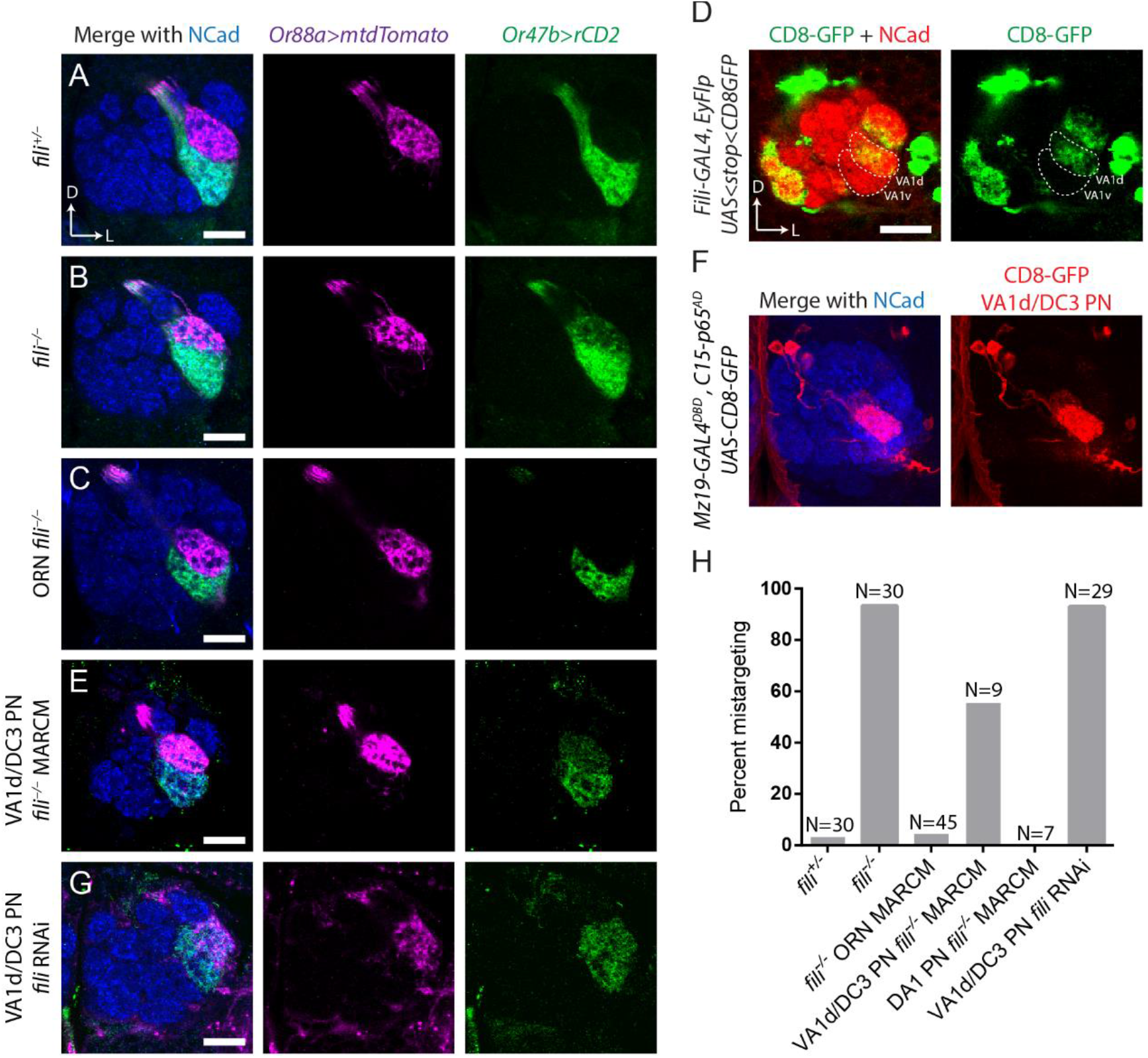
Fili in VA1d/DC3 PNs prevents ectopic targeting of VA1v ORN axons to VA1d glomerulus. (*A*) VA1d and VA1v ORNs, labeled by *Or88a-mTdT* (magenta) and *Or47b-rCD2* (green) respectively, target their axons to two neighboring glomeruli in a stereotyped manner in *fili*^+/−^ animals (mistargeting in 2/30 antennal lobes). (*B*) In *fili*^−/−^ animal, VA1v ORN axons invade VA1d glomeruli (mistargeting in 29/31 antennal lobes). (*C*) Deletion of *fili* in most ORNs using *ey-FLP* MARCM combined with cell-lethal strategy does not alter VA1v ORN axon targeting (mistargeting in 2/45 antennal lobes). (*D*) Expression of Fili in VA1v and VA1d PNs, shown by CD8-GFP (green) using intersection between *GH146-FLP* with *fili-GAL4*. The corresponding glomeruli are outlined by dashed line. (*E*) Deletion of Fili in anterodorsal neuroblast clone (which include VA1d PNs) using *Mz19-GAL4 hsFLP* MARCM causes VA1v ORN axons to mistarget to VA1d glomeruli (mistargeting in 5/9 antennal lobes). (*F*) *Mz19-GAL4^DBD^, C15-p65^AD^* produces functional GAL4 in only VA1d/DC3 PNs to drive *UAS-mCD8-GFP* expression (shown in red). Maximum z-projection is shown. (*G*) *Mz19-GAL4^DBD^, C15-p65^AD^* drives *UAS-fili-RNAi* (VDRC 44532) causes VA1v ORN axons to mistarget to VA1d glomerulus (mistargeting in 27/29 antennal lobes). (*H*) Quantification of VA1v ORN axon mistargeting from panel (*A-C, E and G*). Scale bars, 20 μm.

We hypothesized that similar to our observation in *GMR86C10*-positive PNs (Fig. 4), Fili expressed in PNs targeting to neighboring glomeruli may repel VA1v ORN axons from targeting to inappropriate glomeruli. Our expression analysis using *fili-GAL4* intersected with *GH146-FLP* is consistent this hypothesis: while VA1v PNs did not express *fili*, neighboring glomeruli VA1d, DC3, and DA1 PNs did (Fig. 5*D*). To more directly test our hypothesis, we generated *fili*^−/−^ neuroblast clones visualized by *Mz19-GAL4* (expressed in DA1 PNs that belong to the lateral neuroblast lineage, and VA1d and DC3 PNs that belong to the anterodorsal lineage). At the same time, we labeled VA1d and VA1v ORNs. When the expression of Fili in VA1d/DC3 PNs was eliminated in anterodorsal neuroblast clones, VA1v ORN axons invaded VA1d glomerulus, similar to the whole animal mutant (Fig. 5 *E* and *H*). By contrast, removing *fili* in the lateral neuroblast including DA1 PNs did not cause VA1v ORN axon targeting defects (Fig. 5*H*).

Because the MARCM strategy generates mutants stochastically, it remained possible that we generated *fili*^−/−^ mutants in other cell types that were not labeled by *Mz19-GAL4* to produce this phenotype. We therefore also used RNAi strategy to knockdown *fili* in only VA1d and DC3 PNs. To achieve specific GAL4 expression, we utilized the split-GAL4 strategy (32, 33) to generate new driver lines specific to VA1d and DC3 PNs. We inserted the hemidriver p65^AD^ component after the last coding exons of C15, a transcription factor that is only expressed by PNs derived from the anterodorsal neuroblast (adPNs) (34), and converted *Mz19-GAL4* to *Mz19-GAL4^DBD^* using the HACK strategy (35, 36). When we intersected these two components, expression of functional GAL4 was restricted to *Mz19*+ adPNs (VA1d and DC3) (Fig. 5*F*). Using this newly developed driver line, we drove *fili* RNAi in only VA1d and DC3 PNs, and found that this manipulation recapitulated VA1d mistargeting for VA1v ORN axons as observed in *fili*^−/−^ mutants (Fig. 5 *G* and *H*). Moreover, VA1v ORN axons did not invade the DC3 glomerulus when Fili was removed from VA1d and DC3 PNs by MARCM or RNAi approach. Thus, analogous to VM5 PNs, expression of Fili on the synaptic partner corresponding to the ectopic targeting site is critical for correct axon targeting of VA1v ORN axons. Together, these results suggest that Fili repel neurites of non-matching synaptic partner class to restrict them to the correct target.

## Discussion

Here we have developed a screen for identifying wiring molecules in the ventromedial region of the *Drosophila* antennal lobe. Through an RNAi screen for cell-surface molecules, we discovered that Fili, an LRR-containing transmembrane protein, participates in the assembly of the *Drosophila* olfactory circuit. Detailed expression and genetic analyses revealed that Fili acts as a repellent in ORN for PN, and in PN for ORN, to prevent invasion of neurites into inappropriate glomeruli.

Prior to this study, Fili has been implicated in cell-cell interaction in the detection of mis-specified cells in the wing disc (17). Apoptosis induced by ectopic expression of *Spineless* is regulated by Fili expression. Along with Caps and Trn (37), Fili appears to participate in short-range cell interaction to support cell survival in the wing disc. In the nervous system, LRR-containing proteins have been widely implicated in neural circuit assembly (38). The unique curved structure of LRR combined with exposed β-sheets on the concave side makes it an effective protein-binding motif (39). Furthermore, different LRRs also form distinct structures, permitting interaction with a diverse collection of proteins (40, 41). These unique binding characteristics enable secreted or membrane-associated LRR proteins to play central roles in diverse aspects of nervous system development and function.

Our results suggest that Fili sends a trans-synaptic repulsive signal between ORNs and PNs. Although some LRR are shown to have binding affinity to themselves (42–45), we observed glomeruli innervated by both Fili positive ORN axons and PN dendrites, arguing against a homotypic mechanism for Fili-mediated repulsion. A more probable explanation is that neurons that require Fili for their targeting specificity express a receptor for Fili and are thus repelled by regions with high Fili expression. Future identification of the Fili receptor would further substantiate this hypothesis. Since VA1v ORNs express both Fili and the presumed Fili receptor, Fili does not appear to mediate ORN-ORN repulsion.

Despite Fili’s expression in many classes of ORNs and PNs, we observed that only a small fraction of them requires Fili for correct axon or dendrite targeting. The sparsity of neurons that manifest observable wiring defect in Fili mutant can be explained by the hypothesis that neurons use a combinatorial and redundant coding strategy to specify their connections (11). As we have observed previously, a typical ORN or PN class uses multiple molecules for their axon or dendrite targeting. Our recent single-cell RNAseq data of both PNs and ORNs has revealed that the cell surface landscape between different classes of PNs is usually distinguished by multiple molecules (34, 46). Therefore, when a single wiring molecule is perturbed, the cell surface landscape may still resemble the original neuronal class more closely than other classes, thus permit correct neurite targeting for many neurons. A more comprehensive understanding of the overarching wiring strategies of this circuit will benefit from simultaneous manipulation of multiple wiring molecules that cooperate in specifying the connectivity individual neuronal classes.

## Material and Methods

### Immunostaining

Tissue dissection and immunostaining were performed according to previously described methods (47). Primary antibodies used in this study include rat anti-DNcad (DN-Ex #8; 1:40; DSHB), chicken anti-GFP (1:1000; Aves Labs), rabbit anti-DsRed (1:500; Clontech), mouse anti-rCD2 (OX-34; 1:200; AbD Serotec), mouse nc82 (1:35; Developmental Studies Hybridoma Bank, [DSHB]), anti-HRP conjugated with Cy5 (1:200; Jackson ImmunoResearch), and rat anti-Fili (1:200; custom produced by Thermo Fisher Scientific against a peptide epitope containing Fili residues 684-701 DDEPEHLYERFDHYEYPD). Fili antibody is pre-absorbed by *fili*^−/−^ larval brains to remove non-specific binding. Secondary antibodies raised in goat or donkey against rabbit, mouse, rat, and chicken antisera were used, conjugated to Alexa 405, FITC, 568, or 647 (Jackson Immunoresearch). Confocal images were collected with a Zeiss LSM 780 and processed with Zen software and ImageJ.

### RNAi Screening

The RNAi screen fly was generated as follows: *C155-GAL4* was recombined with *UAS-dcr2* on the X chromosome. *GMR86C10-LexA, LexAop-mtdT, Or98a-mCD8GFP*, and *Or92a-rCD2* were recombined and located on the second chromosome. *UAS-RNAi* males are crossed to this screening line, and the resulting flies were kept at 25 °C for 2 days after egg laying and then transferred to 29 °C to enhance the GAL4/UAS expression system.

### Generation of *fili* Mutant

We generated two gRNA constructs using the BbsI-chiRNA plasmid (48). One gRNA contains a targeting site 5’ to the start codon of *fili* and one 3’ to start of the second coding exon. These two gRNA constructs were co-injected into *Drosophila* embryos with *vas-cas9*, a *yellow* gene (*y*) mutant allele on the X chromosome, and MI02854 containing an exogenous *y* gene on the second chromosome. G_0_ flies were crossed to balancer flies and individuals in F_1_ generation were selected for loss of *y*. Successful events were balanced and confirmed by sequencing.

### Mosaic Analysis

The *hsFLP* MARCM analyses were performed as previously described (30, 49) with slight modifications. *GMR86C10-GAL4* was used for labeling VM5 PNs and *Mz19-GAL4* is used for labeling VA1d, DC3, and DA1 PNs in adult-stage *Drosophila*.

Larvae (24-48 hour after hatching) were heat shocked for 1 hour to obtain neuroblast clones.

### Transgene Generation

To generate enhancer-LexA lines labeling different PNs, including *GMR86C10-LexA*, gateway vector containing enhancer sequences (24) were recombined into the *pBPnlsLexAp65Uw* vector (33) through LR reaction (Invitrogen) and the resulting constructs were injected into *attP2* and *attP40* landing sites by integrase-mediated transgenesis. *Or92a-rCD2* was made by cloning the rat CD2 coding region (50) downstream of Or92a promoter sequence and transgenic animal is made through P-element transformation (51). *C15-p65^AD^* is generated by co-injecting a gRNA (cloned in pU6-BbsI-chiRNA) targeting the end of the last exon and donor sequence containing homology arms, *p65(AD)::Zip*+, and *3XP3-RFP-SV40* (32). *Mz19-Gal4^DBD^* is generated using the HACK strategy (36). We replaced QF2 sequence of pBPGUw-HACK-G4>QF2 with the DNA binding domain of GAL4 cloned from *pBS-KS-attB2-SA(0)-T2A-Gal4DBD-Hsp70 polyA* and injected it to *Mz19-GAL4; vas-Cas9* embryos.

## Acknowledgements

We thank G. Rubin, H. Bellen, T. Lee, the Vienna Stock Center, Bloomington Stock Center, *Drosophila* Genomics Resource Center, and Addgene for reagents. We thank T. Li, C.N. McLaughlin, A. Shuster, J. Ren, J. Lui, D. Pederick, A. Ward for helpful discussions and comments on the manuscript. This work was supported by NIH Grant R01 DC005982 (to L.L.). L.L. is a Howard Hughes Medical Institute investigator.

## Supplementary Information

**Figure S1.**
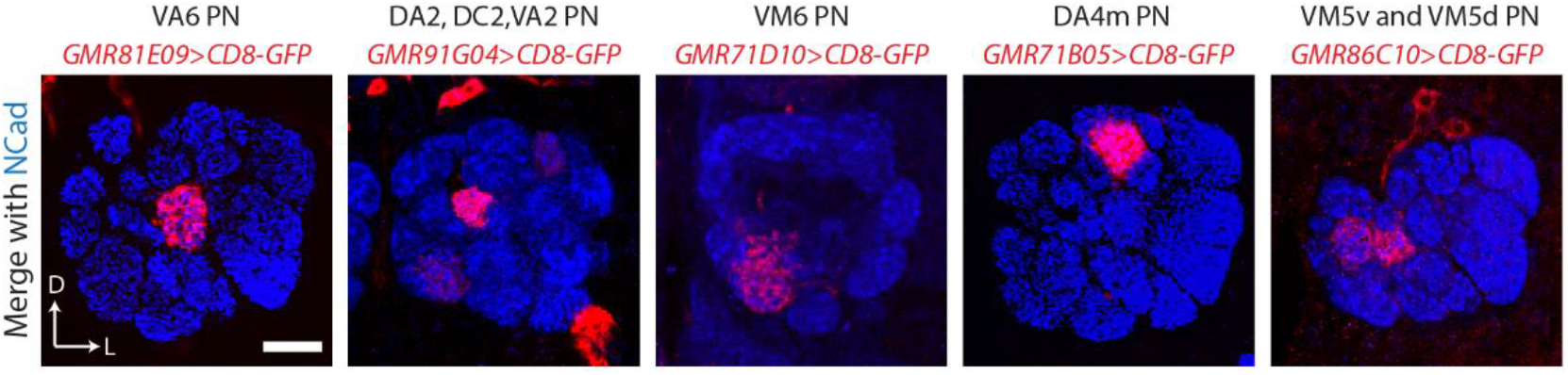
*Enhancer-LexA* lines label specific PN neurons in adult. Confocal sections of adult antennal lobe are shown. Neuronal processes are labeled by five different *enhancer-LexA>LexAop-mCD8GFP* (red) lines. Blue channel shows neuropil staining by the Ncad antibody. Scale bars, 20 μm.

**Figure S2.**
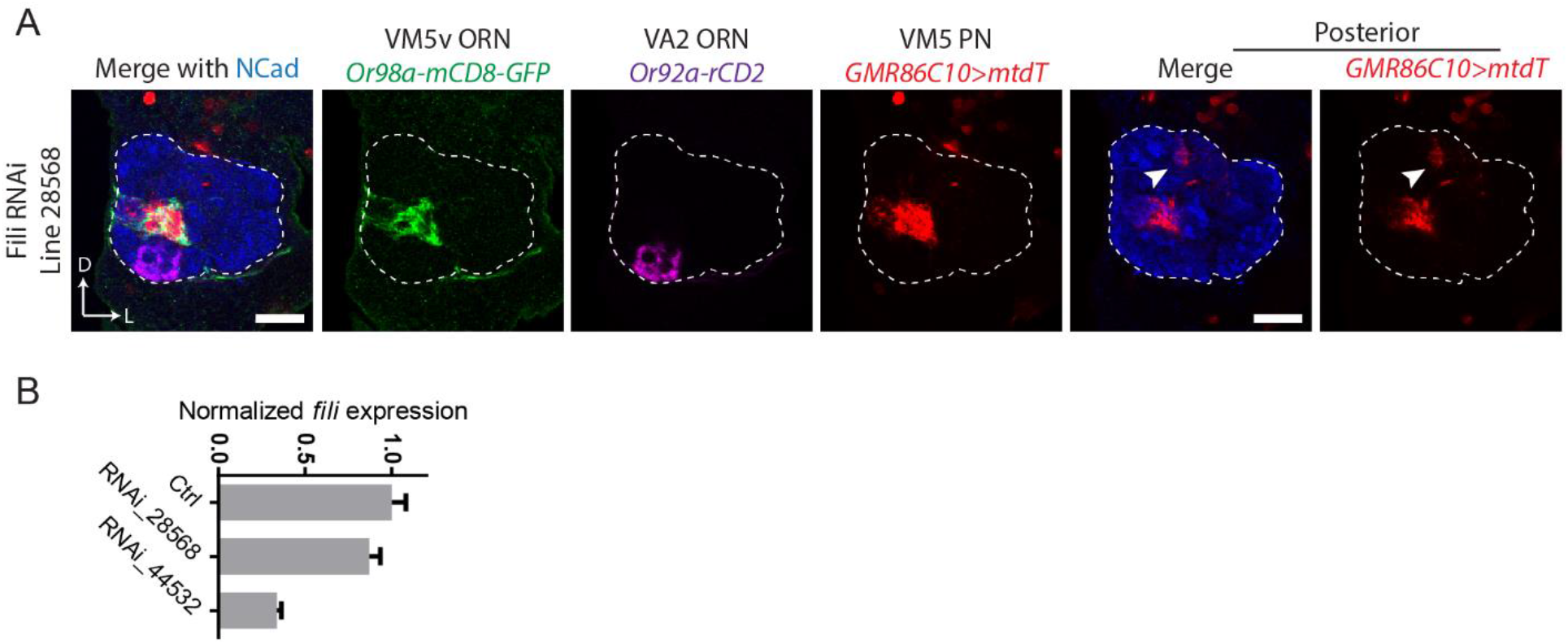
*fili* knockdown by two separate RNAi lines shows PN dendrite targeting defect. (*A*) *C155-GAL4* drives *UAS-fili-RNAi* (Bloomington 28568) shows PN dendrite targeting defect. Ectopic PN target are indicated by arrowhead on the posterior section. (mistargeting in 7/20 antennal lobes). (*B*) Quantitative PCR (qPCR) measurement of the knockdown efficiency of *fili* using two *UAS-fili-RNAi* lines. *C155-GAL4* was crossed with either *w^1118^* (control) or two Fili-RNAi lines. mRNA was extracted from 5-day-old adult fly heads (n=3 biological replicates of 10 heads pooled for each condition). Expression levels are normalized to *actin5C*. Error bars show SEM. Scale bars, 20 μm.

**Figure S3.**
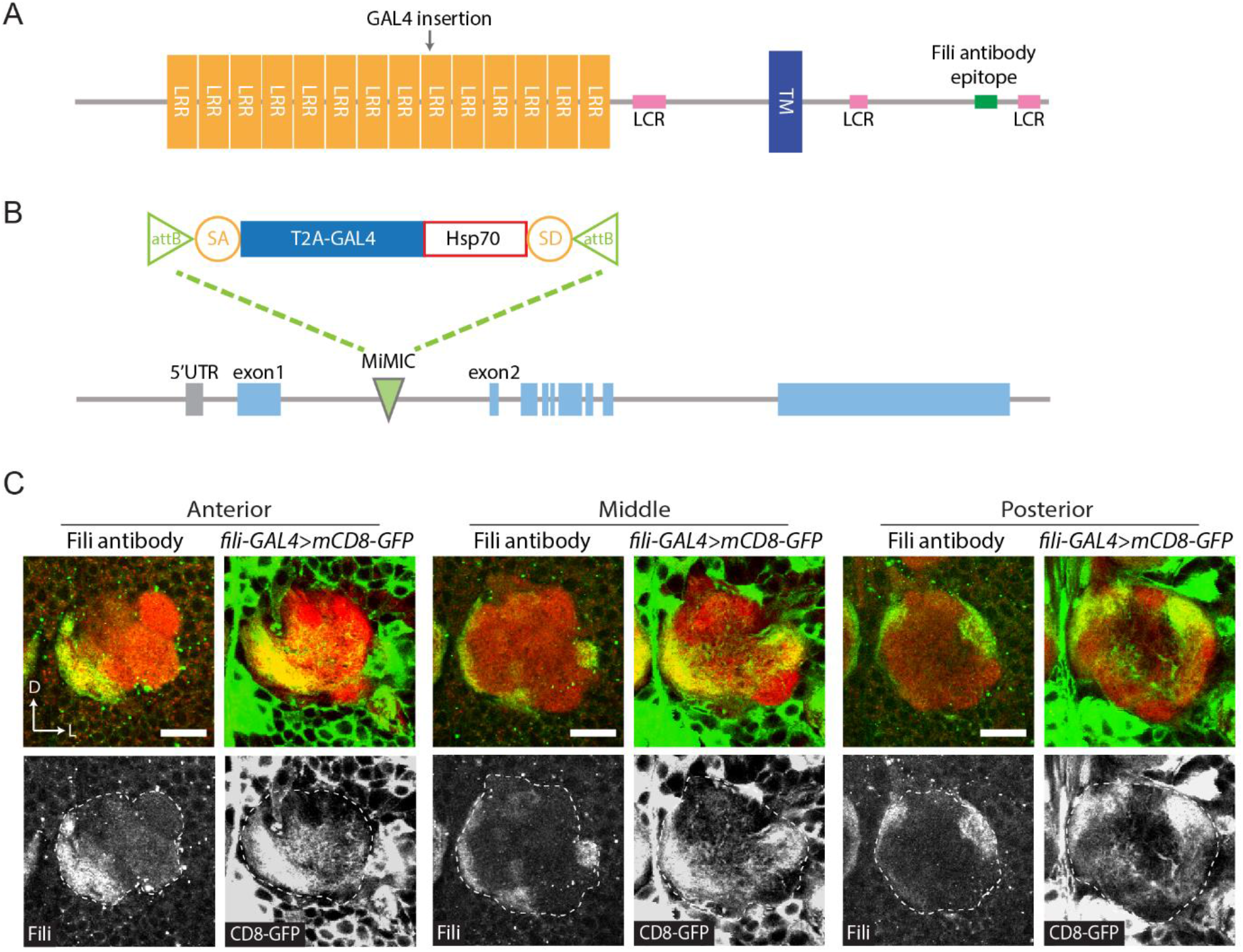
Fili expression revealed by antibody and transcriptional reporter GAL4 showed similar pattern. (*A*) Fili protein domain prediction. The blue bar (TM) represents transmembrane domain. Orange bars (LRR) indicates leucine-rich repeat domains. Pink bars (LCR) indicate low-complexity regions. The epitope for Fili antibody is represented by the green bar. (*B*) Design of *fili-GAL4*. A *T2A-GAL4* cassette was inserted into a MiMIC locus in the first intron of *fili*. SA, splicing acceptor. Hsp70, terminator sequence of *Hsp70*. SD, splicing donor. Blue bars: coding exons of *fili*. (*C*) Fili antibody staining and *fili-GAL4* shows consistent pattern at 48h APF. Three optical sections (anterior, middle, and posterior) are shown for both Fili antibody staining and CD8-GFP (green). Neuropil is visualized by antibody against either NC82 or NCad (red). Dashed lines outline the antennal lobe. Scale bars, 20 μm.

**Figure S4.**
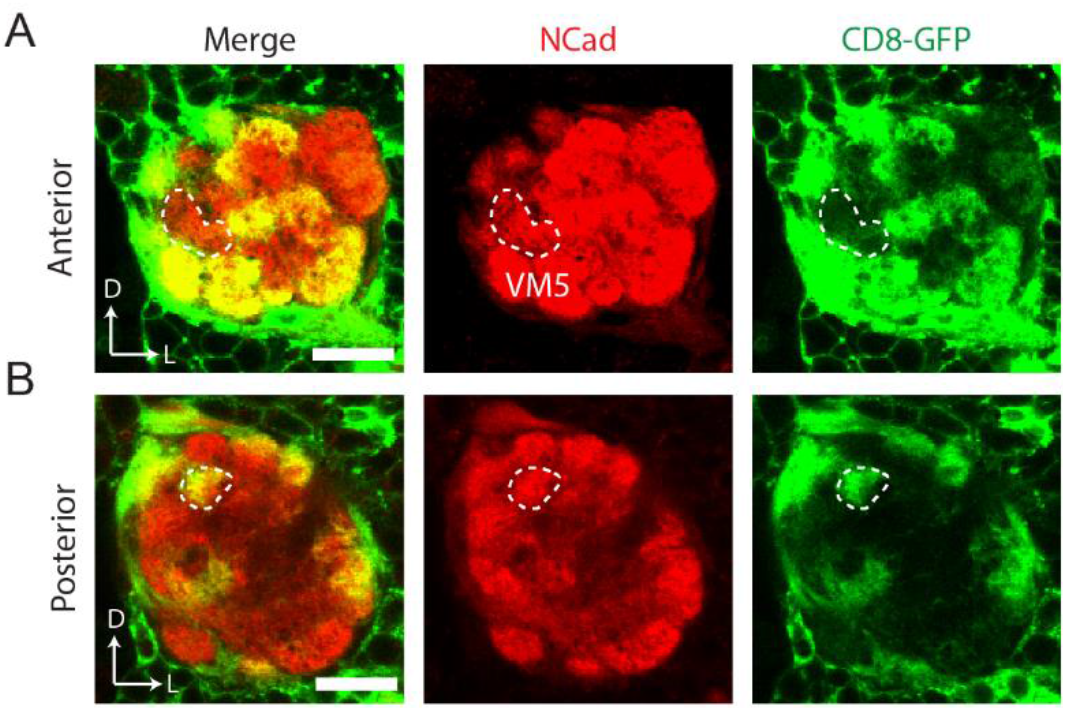
The ectopic target site of VM5 PN dendrites is innervated by *fili-GAL4*+ ORN axons. *ey-FLP* intersecting with *fili-GAL4* together with *UAS-FRT-stop-FRT-mCD8GFP* shows *fili* expression pattern in ORNs in the 48 hours APF antennal lobe. (*A*) VM5v and VM5d glomeruli (outlined by dashed line) do not have detectable ORN Fili signal. (*B*) The ectopic targeting site (outlined by dashed line) of VM5 PNs have high ORN Fili signal. Scale bars, 20 μm.

**Figure S5.**
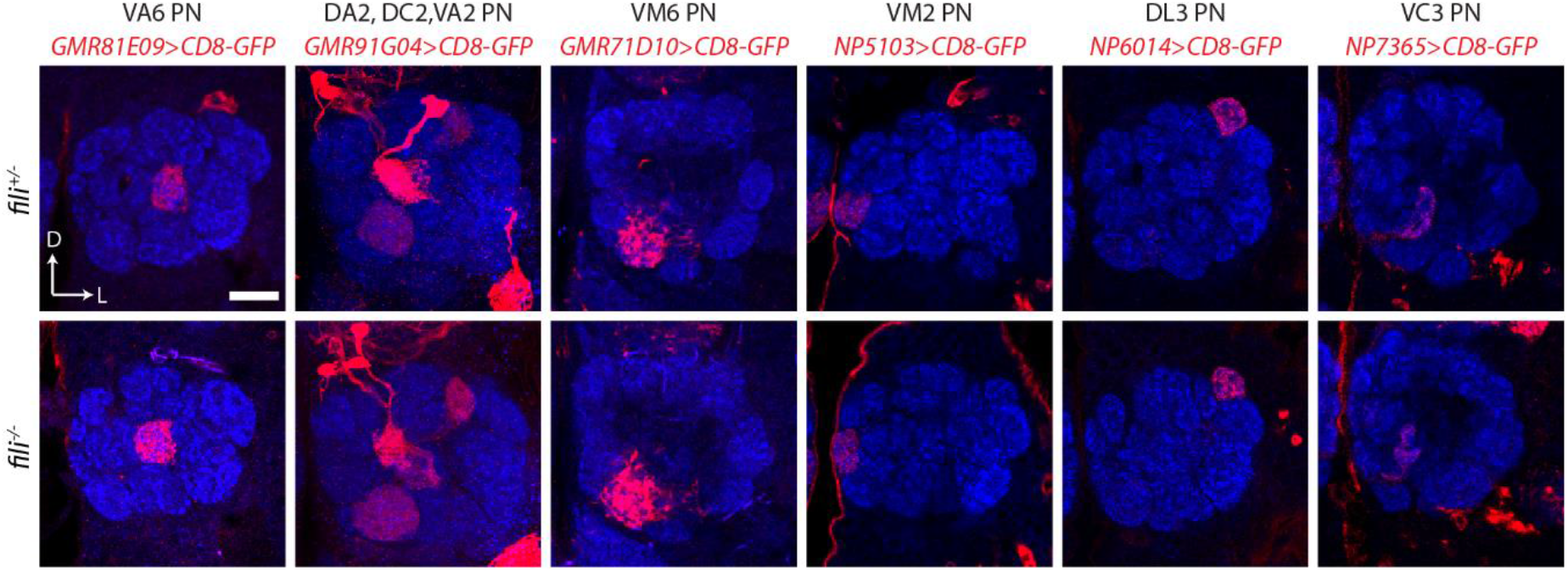
Fili is not required for the targeting of the 8 tested PN classes. PN dendrites (red) are visualized in *fili*^+/−^ control animal and *fili*^−/−^ for VA6 PN (*GMR81E09-GAL4*); DA2, DC2, and VA2 PN (*GMR91G04-GAL4*); VM6 PN (*GMR71D10-GAL4*); VM2 PN (*NP5103-GAL4*); DL3 PN (*NP6014-GAL4*); VC3 PN (*NP7365-GAL4*). Maximum z-projection is used for *GMR91E04-GAL4* images to show dendrite targeting of all 3 PN classes, all other images are single sections. Scale bar, 20 μm.

**Figure S6.**
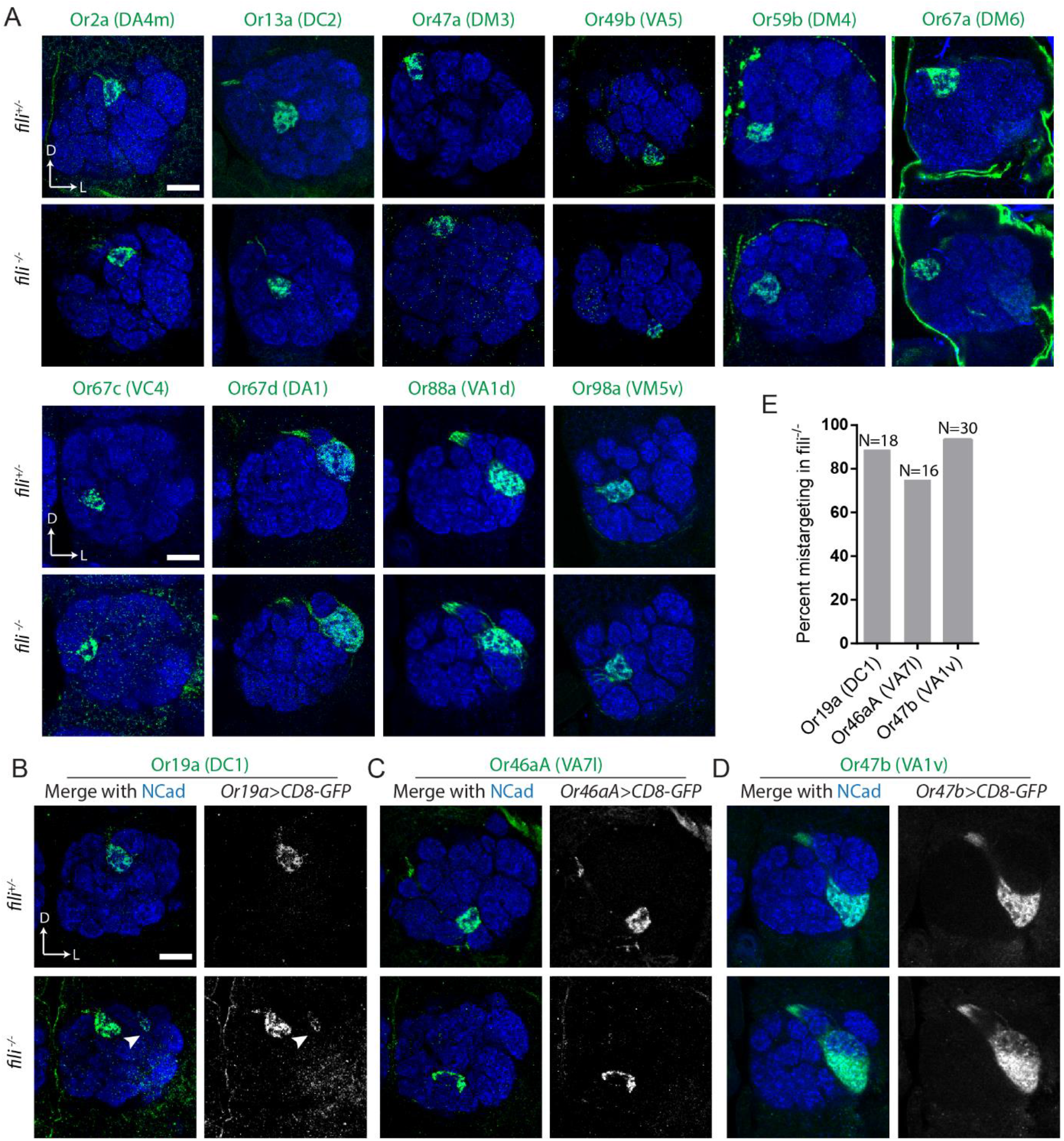
Fili is required for the axon targeting of some ORN classes. ORN axons (green) in *fili*^+/−^ and *fili*^−/−^ animals visualized using either Or-promoter fused upstream of GAL4 to drive *UAS-mCD8-GFP* expression, or direct fusion of Or-promoter and *mCD8-GFP, mtdTomato*, or *rCD2*. (*A*) ORN classes whose axon targeting does not require Fili. (*B*) *Or19a-mCD8-GFP* labeled DC1 ORN axons shows an ectopic target on the lateral side of DC1 in *fili*^−/−^. Arrowhead points to the ectopic target. (mistargeting in 16/18 antennal lobes). (*C*) *Or46a-mCD8-GFP* labeled VA7l ORN axons are misshapen in *fili*^−/−^ animal but no ectopic target site is observed (mistargeting in 12/16 antennal lobes). (*D*) Or47b-rCD2 labeled VA1v ORN axons invade the VA1d glomeruli in *fili*^−/−^ animals (mistargeting in 28/30 antennal lobes). (*E*) Quantification of mistargeting of DC1, VA7l, and VA1v ORN axons shown in (*B–D*). Scale bars, 20 μm.

